# *Candidatus Ornithobacterium hominis* sp. nov.: insights gained from draft genomes obtained from nasopharyngeal swabs

**DOI:** 10.1101/326074

**Authors:** Susannah J Salter, Paul Scott, Andrew J Page, Alan Tracey, Marcus C de Goffau, Bernardo Ochoa-Montaño, Clare L Ling, Jiraporn Tangmanakit, Paul Turner, Julian Parkhill

## Abstract

*Candidatus Ornithobacterium hominis* sp. nov. represents a new member of the Flavobacteriaceae detected in 16S rRNA gene surveys from Southeast Asia, Africa and Australia. It frequently colonises the infant nasopharynx at high proportional abundance, and we demonstrate its presence in 42% of nasopharyngeal swabs from 12 month old children in the Maela refugee camp in Thailand. The species, a Gram negative bacillus, has not yet been cultured but the cells can be identified in mixed samples by fluorescent hybridisation. Here we report seven genomes assembled from metagenomic data, two to improved draft standard. The genomes are approximately 1.9Mb, sharing 62% average amino acid identity with the only other member of the genus, the bird pathogen *Ornithobacterium rhinotracheale.* The draft genomes encode multiple antibiotic resistance genes, competition factors, *Flavobacterium johnsoniae*-like gliding motility genes and a homolog of the *Pasteurella multocida* mitogenic toxin. Intra- and inter-host genome comparison suggests that colonisation with this bacterium is both persistent and strain exclusive.

## AUTHOR SUMMARY

The nasopharynx is part of the respiratory tract and hosts a unique microbial community that is established during infancy, changing throughout life. The nasopharyngeal microbiome is important to study as it includes bacteria that can cause diseases such as otitis media or pneumonia, as well as non-pathogenic species. In Maela, a refugee camp in Thailand, we identified a prevalent bacterial species colonising children under the age of two years and occasionally their mothers. We were not able to culture it from frozen swabs, but could visualise the cells microscopically using a fluorescent probe. Its genetic signature can be seen in published data from several countries suggesting that the species may be widespread. From analysis of the genome we confirm it is highly divergent from its closest characterised relative, the respiratory pathogen *Ornithobacterium rhinotracheale*, which infects turkeys, chickens and other birds. We propose the name *Candidatus Ornithobacterium hominis* sp. nov.

## INTRODUCTION

During previous work on the nasopharyngeal microbiota of children in the Maela refugee camp in Thailand, an abundant unclassified taxon was discovered through 16S rRNA gene sequencing [1]. It was >99% identical to other unclassified sequences reported in nasopharyngeal samples from the Gambia [2, 3], Kenya [4], and Australia [5], and the gene shared 93% nucleotide identity with that of the avian respiratory pathogen *Ornithobacterium rhinotracheale* (ORT). On the basis of 16S rRNA gene similarity the taxon was presumed to represent a new species of Flavobacteriaceae, closely related to the genus *Ornithobacterium.* The taxon is of interest because it was ubiquitous in the study group of 21 children, appearing to be a persistent coloniser and at a proportional abundance up to 71%. The 16S rRNA gene sequences could be divided into three oligotypes [6]: each appeared to be carried persistently and exclusively by their host [1].

As the bacterium could not be cultured from swabs, ten DNA samples from the initial study were selected for metagenomic sequencing to maximise recovery of the genome of interest while representing a range of children, ages, and 16S rRNA gene oligotypes: seven were successfully sequenced. The extracted genomes were then used to design a PCR-based prevalence screen for samples from the Maela cohort and a fluorescent probe to visualise the cells in mixed samples. On the basis of this genomic analysis, we propose the unclassified taxon as *Candidatus Ornithobacterium hominis* sp. nov. (OH).

## RESULTS

Genomes of OH were assembled from metagenomic data generated on an Illumina Miseq. Despite significant loss of sequence coverage to human and other bacterial genomes, two samples assembled into 9 and 15 contigs from OH, yielding draft genomes predicted to be nearly complete based on the detection of all ribosomal protein genes and by inter-sample comparison. A further five samples assembled into larger numbers of smaller contigs, which were aligned to the draft genomes and found to cover most of the expected genome (Table 1). The OH genome is approximately 1.9Mb, 20% smaller than its closest relative ORT.

**Table 1:** Sequenced sample information; well-assembled draft genomes are highlighted in green.

| Child | Age (months) | Sample | Date | Genome ID (accession #) | No. extracted contigs | Total length | Max contig length | Proportional abundance in sample [1] |
| --- | --- | --- | --- | --- | --- | --- | --- | --- |
| ARI0073 | 11 | 08B08946 | Nov 2008 | OH-22797 (ERS2321173) | 302 | 1.81Mb | 32kb | 56% |
|  | 15 | 09B02844 | Mar 2009 | OH-22872 (ERS2321176) | 41 | 1.93Mb | 509kb | 38% |
|  | 21 | 09B09452 | Sep 2009 | OH-22767 (ERS2321172) | 15 | 1.93Mb | 611kb | 31% |
| ARI0106 | 20 | 09B08231 | Aug 2009 | OH-22298 (ERS2321170) | 183 | 1.80Mb | 68kb | 67% |
| ARI0218 | 14 | 09B00559 | Jan 2009 | OH-22819 (ERS2321175) | 172 | 1.42Mb | 51kb | 49% |
|  | 19 | 09B06140 | Jun 2009 | OH-22803 (ERS2321174) | 9 | 1.87Mb | 684kb | 62% |
| ARI0484 | 15 | 09B08221 | Aug 2009 | OH-22763 (ERS2321171) | 205 | 1.88Mb | 57kb | 51% |

## Prevalence

An OH specific real-time PCR detection protocol for V2-V5 of the 16S rRNA gene was designed using full length gene sequences and tested on archived nasopharyngeal swabs (NPS) in STGG (skim milk, tryptone, glucose and glycerol storage medium) and on metagenomic DNA of known bacterial composition. This PCR screen was then applied directly to STGG from archived NPS of 100 randomly selected 12-month old infants in Maela, and concurrent swabs from their mothers. A second PCR screen was developed targeting the toxin gene *toxA* and was also performed on the archived NPS. The two PCR targets were concordant in infant samples, resulting in 42 positive and 58 negative results, giving an estimated carriage prevalence among 12 month old infants in Maela of 42% (95% CI: 32.3-51.7). From the mothers 2 samples were positive, 93 negative, and 5 were either equivocal or nonconcordant (Table 2). The cycle threshold (Ct) values were higher in maternal samples than infants, which may indicate a lower bacterial load.

**Table 2:**
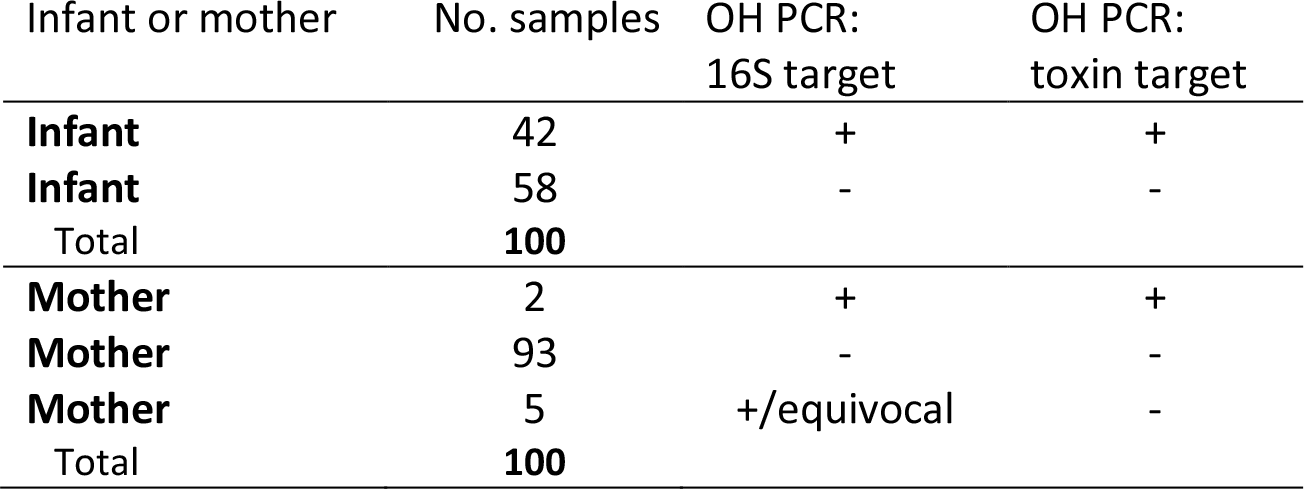
OH prevalence in mother and infant samples.

## Similarity to *O. rhinotracheale*

The draft genomes appear to be very distantly related to ORT by several measures, summarised in Table 3. The average nucleotide identity (ANI) between OH-22767, OH-22803 and ORT UMN-88 [7] can only be calculated from a small fraction of the genome. The two-way average amino acid identity (AAI) between the two drafts and UMN-88 is approximately 62% based on three quarters of predicted proteins. Another measure, the reciprocal percentage of conserved proteins (POCP) [8], may be used to gauge the relatedness of two genomes at the genus level. To be considered conserved for this measure a gene must share >40% amino acid identity over >50% of its length: two members of the same genus are expected to have at least half of their proteins in common. The POCP between UMN-88 and the draft genomes is approximately 58%. Although these figures are based on incomplete genomes, as 50.7% of UMN-88 proteins are conserved in OH from these data, they are likely to be very distantly related members of the same genus.

**Table 3:** Comparison of OH draft genomes with ORT genome UMN-88.

| Draft genome | Average nucleotide identity (ANI) |  | Average amino acid identity (AAI) |  | Percentage conserved proteins (POCP) |  |
| --- | --- | --- | --- | --- | --- | --- |
| **OH-22767** | 81.7% | 7.2% genome aligned | 61.8% | 75.0% proteins aligned | 57.6% | 50.7% ORT conserved<br>66.7% OH conserved |
| **OH-22803** | 77.7% | 5.8% genome aligned | 61.5% | 77.7% proteins aligned | 58.1% | 50.7% ORT conserved<br>68.2% OH conserved |

## Detection in swab material using fluorescent hybridisation

A fluorescent probe targeting the V5 region of the 16S ribosomal RNA was designed for OH using full-length sequences of the 16S rRNA gene. It was used to visualise OH cells directly from nasopharyngeal swab samples. OH is a Gram negative bacillus, often observed in pairs and occasionally longer chains, similar to the morphology of ORT (Figure 1). ORT was tested in parallel and could not be visualised with the same fluorescent probe.

**Figure 1:**
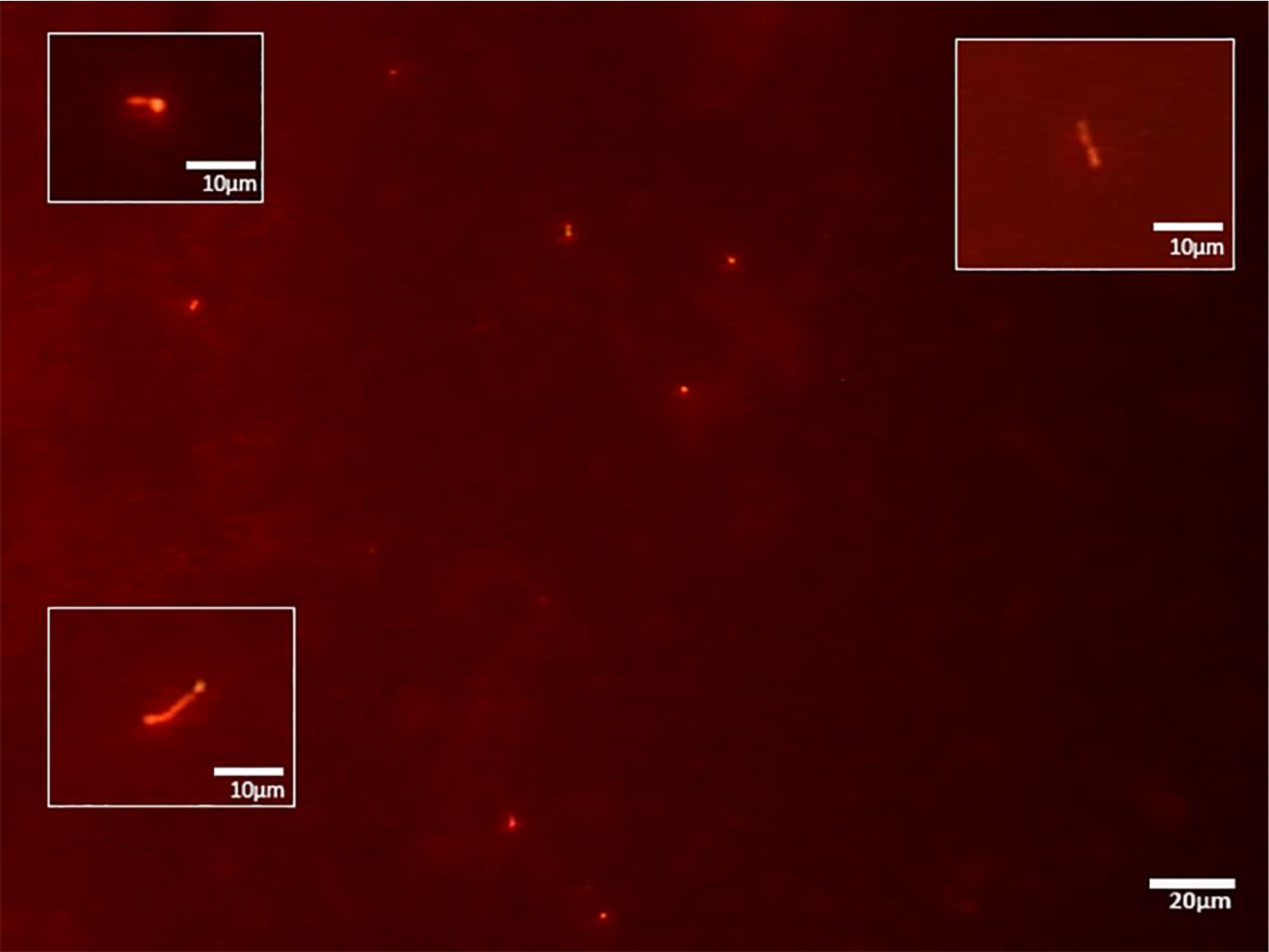
OH cells visualised with fluorescent ribosomal probe.

## Core genome

The core genes shared between the draft genomes and lower quality assemblies, along with the accessory genes unique to each, were calculated using Roary [9]. Due to the low quality assemblies containing gaps, just 935kb or approximately 50% of the draft genome size was identified as “core genome” using this analysis. A core genome phylogeny was generated using >13,000 SNPs identified in this shared sequence [10]. Samples taken from the same child at different dates are very similar (Figure 2), adding to the initial 16S rRNA gene oligotype data that inferred long-term carriage of the same or closely related strains in this cohort. Features of interest that are present in the core genome include a large toxin gene *toxA* and gliding motility-associated genes.

**Figure 2:**
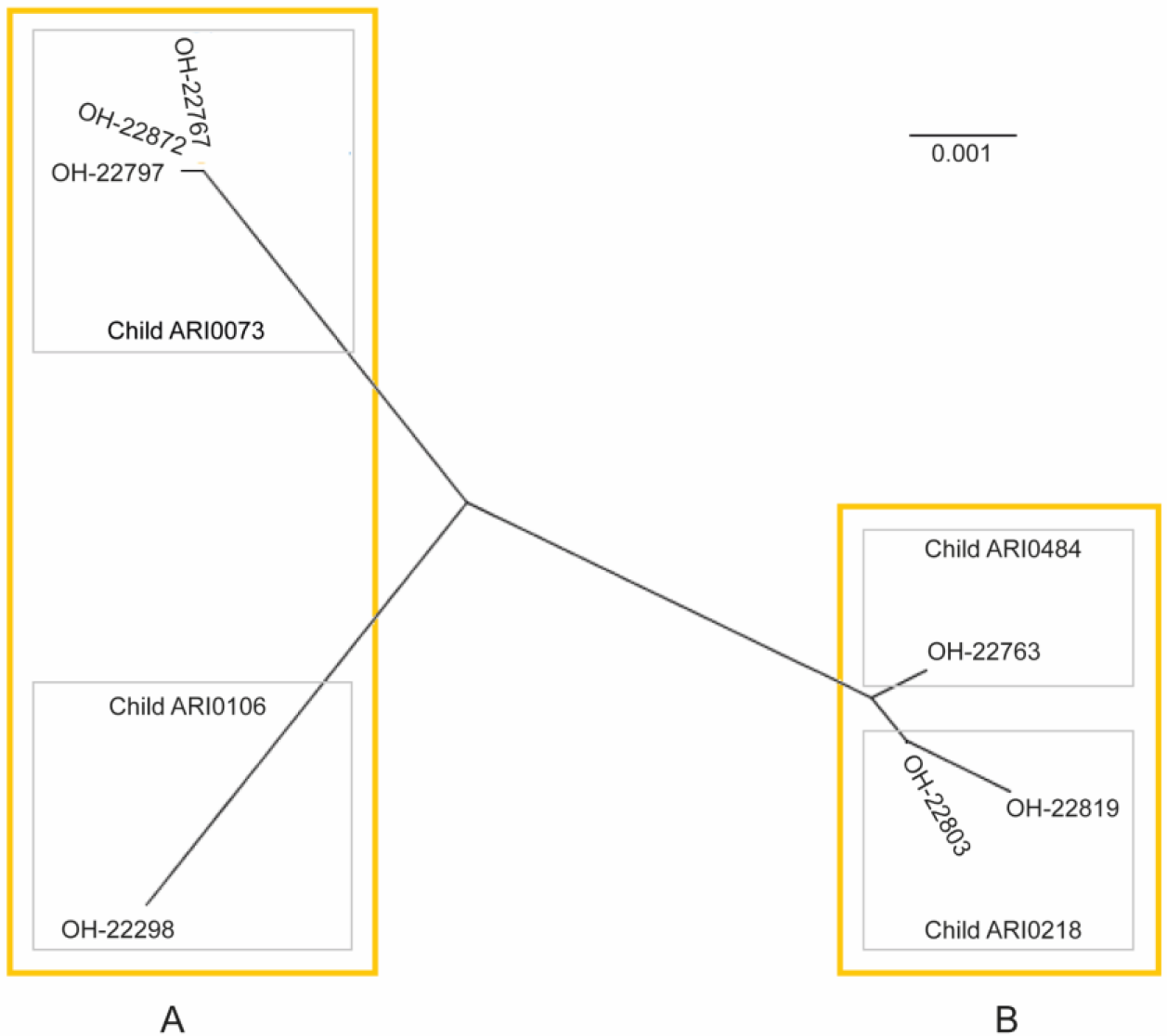
Unrooted phylogenetic tree built from 935kb “core genome” sequence from high and low quality assemblies using RAxML. Boxes represent groups based on accessory genome discussed below.

The 3.8kb gene *toxA* is present in all sequenced samples and was detected by PCR in all 16S-positive swabs from children. It is predicted to produce a secreted toxin similar to the *Pasteurella* mitogenic toxin (PMT). Although OH ToxA and PMT share only 35% amino acid identity overall, there is greater conservation around the predicted active sites, and the modelled structure of the C-terminal domain is extremely similar (Figure 3).

**Figure 3:**
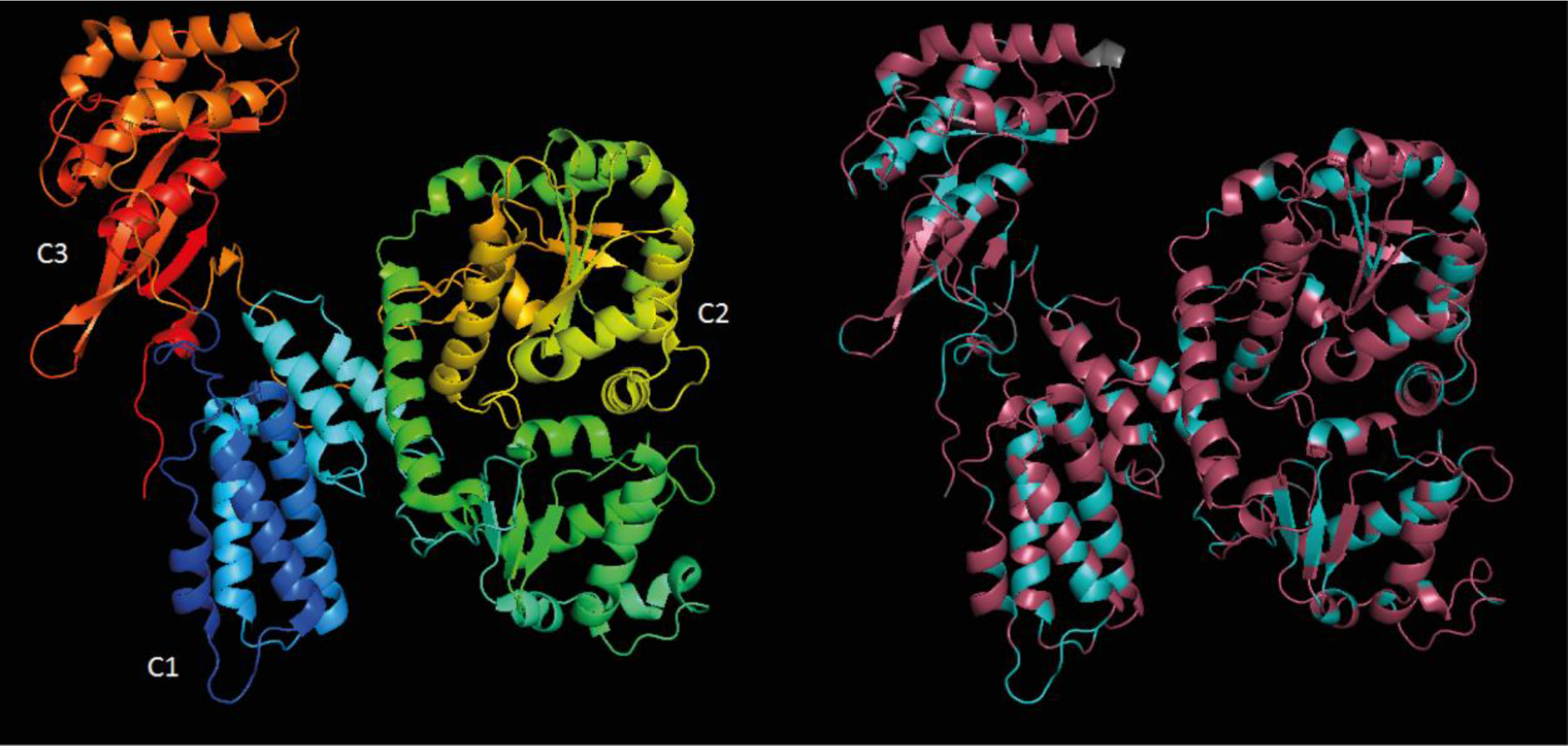
Modelled structure of *Pasteurella* mitogenic toxin (PMT) C-terminal domain, left, and of equivalent region of OH ToxA, right. The PMT model is coloured on a rainbow spectrum to indicate position. The OH model is coloured according to amino acid identity to PMT with identical residues in blue, non-identical in purple, and insertions in grey. In PMT the C1 region is responsible for plasma membrane localisation, the C3 region forms an active pocket and is responsible for mitogenic activity in mammalian cells.

The core genome includes a full complement of 14 *gld* genes that are homologs of those required for gliding motility by *Flavobacterium johnsoniae*. This mechanism involves the movement of an adhesin around the cell membrane in a helical path, thereby pulling the bacterium rapidly along a substrate [11]. Most of these genes in *F. johnsoniae* are also components of the Bacteroidetes type IX secretion system (T9SS) [12, 13], which is responsible for secretion of *F. johnsoniae* gliding adhesins SprB and RemA. However *sprB* has not been identified in OH.

## Accessory genome

Around half of the accessory genome is made up of hypothetical protein genes, most of which are similar to those of other members of the Flavobacteriaceae such as ORT, *Capnocytophaga, Chryseobacterium, Elizabethkingia, Flavobacterium, Riemerella,* or *Weeksella*. It also includes evidence of mobile elements and associated drug resistance genes, Rhs (rearrangement hotspot) genes, and two distinct lipopolysaccharide production clusters.

A portion of the hypothetical protein genes encode the *Fibrobacter succinogenes* major paralogous domain (PF09603). Up to 17 of these genes are present in each genome. A quarter of them are also predicted to possess an immunoglobulin-like fold (IPR013783). No two samples have a complement of completely identical predicted genes, but more are shared between samples that group together based on the core genome (Figure 2).

The genomes contain different complements of bacteriophage associated genes, Type I, II and III restriction modification systems, diverse variants of DNA degradation genes *dndCDE* [14], abortive infection systems, transfer and mobilisation genes. In some cases these are present in small regions of <10kb and lack any clear structure, while in others they are part of a defined element such as a 30kb tailed bacteriophage (22803_00899-00942), or a 19kb element (22803_01685-01710) containing efflux genes and flanked by 350bp imperfect direct repeats. All sequenced samples possess efflux transporters of the MATE and RND families, but genomes from cluster A (Figure 2) have an additional B1 metallo-beta lactamase (22767_01182), streptogramin lyase (22767_01181) and *ampC* gene (22767_01179) within a partially-assembled mobile element. Genomes from cluster B possess an extended spectrum beta lactamase (ESBL) gene *per1* on the well-characterised transposon Tn4555 [15].

The accessory genome of strains from cluster A encodes a number of elements containing the conserved RHS repeat-associated core domain (TIGR03696) with extremely variable C termini and unique hypothetical protein genes immediately downstream. There is one large Rhs gene of >9kb, 22767_01758, that encodes a protein sharing signatures with the *Salmonella* plasmid virulence protein SpvB (IPR003284) and the bacterial insecticide toxin TcdB (IPR022045), while the other smaller ORFs may be the dissociated tips generated from lateral acquisition of variable C termini as seen in *Serratia marcescens* [16]. This large Rhs gene with displaced tips is only found in samples from two children, ARI0106 and ARI0073: some but not all of the Rhs gene tips differ between them. These samples also possess a further Rhs gene with no displaced tips, that has two predicted phospholipase D domains (IPR001736). Some Rhs proteins have been shown to act as competition factors [17], consistent with OH being a persistent member of the nasopharyngeal microbiota.

Although all genomes possess a 30kb lipopolysaccharide (LPS) production gene cluster, 11kb of this region differs between samples from clusters A and B, containing a different complement of transferases and synthases, and suggesting the presence of at least two serotypes in the species. The A variant (22767_00486-00496) includes an ABC transporter and several transferase and synthase genes 45-55% identical to those of ORT, while the B variant (22803_00491-00502) has a Wzx flippase and genes that are more similar to *Chryseobacterium* or other Flavobacteriaceae.

## DISCUSSION

Previously there has been little evidence available for OH colonising adults in Thailand or other countries, as most nasopharyngeal microbiota studies target young children. By screening 100 pairs of mother/child samples (Table 2), 2% of samples from mothers and 42% from infants were unequivocally PCR-positive for OH. The prevalence at 12 months of age could be estimated as 32.3-51.7% (95% CI) using this screen but the maternal prevalence could not be accurately measured from this number of samples. The Ct values for mothers may indicate a lower bacterial load, leading to a modest under-detection of colonisation by PCR. It contrasts with other bacteria such as *Streptococcus pneumoniae* which is commonly carried in both adults and infants in Maela: among these 100 pairs of swabs 31% from mothers and 79% from infants were culture positive for *S. pneumoniae*.

Preliminary efforts have been made to culture the bacterium from nasopharyngeal swabs expected to contain a large proportion of this species, following protocols appropriate for ORT or various nasopharyngeal bacteria. OH has not yet been successfully cultured from any archived sample. Furthermore, the frequency of fluorescent cells observed during ribosomal probing of mixed samples was not as high as expected from the 16S analysis. This difficulty culturing the bacterium may be due to unknown metabolic requirements, or it may be that the cells have not survived storage due to environmental stresses or lysis.

In earlier work [1] it was noted that each child was colonised with only one of the three detected 16S rRNA gene oligotypes representing OH. Colonisation was persistent, i.e. constituting >5% of the proportional abundance of taxa for at least 5 consecutive months, in 13 out of 21 children but was detected in all children at some point during the study. Here we describe multiple similar OH genomes taken from time-points that are 4-10 months apart in two children, adding evidence to the hypothesis that long-term colonisation is restricted to a particular strain for each host. Given the high prevalence of OH in Maela (42% of 12 month old children) and the genetic diversity observed between contemporaneous samples from only four hosts, this exclusion of diversity from the host may be explained by microbial competition systems such as the SdpABC-like toxin system identified in OH-22298 [18] or Rhs proteins [17]. Due to the high frequency of clinical pneumonia in Maela (0.73 episodes per child year [19]), the children and their microbiota are frequently exposed to beta lactam antibiotics. In all sequenced samples we found evidence of horizontally acquired drug resistance genes, which may also aid persistent colonisation.

Bacterial LPS may confer advantages in adhesion and avoidance of complement mediated cell lysis, although it is also a key target for the host immune system [20]. The gene content of LPS cluster variant A is somewhat similar to that of the ORT serotype A (Figure 4), the most common of the 18 known serotypes [21] and the only ORT LPS type currently represented in public sequence databases. Despite individual gene similarity to LPS clusters of several *Chryseobacterium* species including *C. gallinarum* and *C. senegalense,* OH variant B in its entirety does not resemble that from any known genomes.

**Figure 4:**
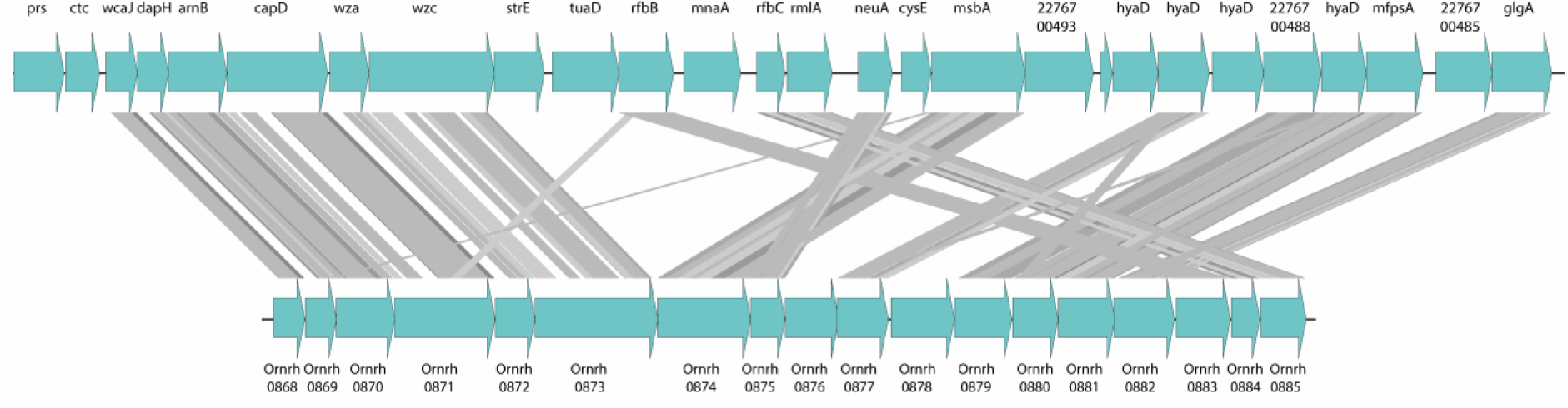
tblastx alignment of OH LPS variant A (OH-22767, top) and ORT serotype A (type strain DSM 15997, bottom).

Gliding motility is often associated with firm dry surfaces [22] though it is also found among oral bacteria such as those from the genera *Cytophaga* and *Capnocytophaga.* The ability to move independently in the environment is advantageous for scavenging nutrients, for complex biofilm formation, and to bring about contact with other bacterial or host cells. The *gld* genes that are required for gliding motility among the Flavobacteriia, Cytophagia and Sphingobacteriia overlap with those of the T9SS, so a subset of these genes are also found in non-motile relatives [23]. Despite possessing all 14 *gld* genes and further required genes *sprAET,* these OH genomes do not include an SprB-like adhesin and putative gliding motility must be confirmed phenotypically.

PMT is a toxin produced by some serovars of *Pasteurella multocida* that causes a range of host pathologies including nasal bone resorption [24], lower respiratory tract disease [25, 26], and dermonecrotic wound infections [27], and has also been shown experimentally to affect the heart [28], liver [29], and bladder [30]. It acts by deamidating the α subunits of several heterotrimeric G proteins, activating mitogenic signalling pathways [31, 32]. In the Maela cohort there are no reports of PMT-like toxin mediated disease, strongly suggesting that despite structural similarities the expression or function of OH ToxA is different to that of *P. multocida.*

In conclusion, we have assembled seven genomes representing a new species of nasopharyngeal bacteria, proposed as *Candidatus Ornithobacterium hominis* sp. nov., from nasopharyngeal swabs collected from a cohort of children in the Maela refugee camp in Thailand. Although phenotypic characterisation is not yet possible due to undetermined storage or culture requirements, several points of interest have been identified for further investigation. These include the predicted gliding motility phenotype, two lipopolysaccharide variants, and the production of a protein similar to the *Pasteurella* mitogenic toxin. The prevalence of OH colonisation appears to be approximately 42% of 12 month old children in Maela refugee camp and needs to be estimated in other populations.

## MATERIAL AND METHODS

### Samples

Between 2007 and 2010, a cohort of 955 infants born in the Maela refugee camp on the Thailand-Myanmar border were followed from birth until 24 months of age in a study of pneumococcal colonisation and pneumonia epidemiology [19, 33]. Nasopharyngeal samples were collected from each infant at monthly intervals using Dacron tipped swabs (Medical Wire & Equipment, Corsham, UK). Immediately following collection, the NPS were placed into STGG transport medium and frozen at -80°C within 8 hours. An additional nasopharyngeal swab was taken if the infant was diagnosed with pneumonia according to WHO clinical criteria [34].

### Ethics statement

Infants were eligible for inclusion in the cohort study if informed written consent had been obtained from the mother during the antenatal period. Participants and their data were anonymised using a 4-digit code prefixed by “ARI-”. The study protocol was reviewed and approved by the ethics committees of the Faculty of Tropical Medicine, Mahidol University, Thailand (MUTM-2009-306) and University of Oxford, UK (OXTREC-031-06).

### Prevalence screen

Prevalence of carriage in the infant population of Maela was estimated using a qPCR screen direct from NPS storage medium. The 12-month routine samples from 100 randomly selected infants (excluding twins) were used. The age of child at time of sampling ranged from 359-377 days, median 365 days. Concurrent swabs were also acquired from the mothers and screened to assess maternal carriage in relation to infant carriage. qPCR was performed on an Applied Biosystems 7500 RT PCR machine using PowerUp SYBR Green mastermix (Applied Biosystems) in 20μl reaction volumes. The 16S rRNA gene screen targets the V2-V5 region with forward primer CTTATCGGGAGGATAGCCCG and reverse GAAGTTCTTCACCCCGAAAACG, yielding a 700bp product under the conditions: 94°C for 5 minutes (cell lysis), then 40 cycles of 94°C for 30 seconds, 53°C for 30 seconds, 68°C for 1 minute, ending with a melt curve. A positive result was a Ct<40 and peak melting temperature (Tm) of 80-86°C. The ToxA gene screen with forward primer TATCTCTCACAGAGCTAGGCTTGAGCGTGG and reverse TGCTATATTTGGGAAAGGCGCATGAATACC yields a 1.95kb product under the conditions: 94°C for 5 minutes (cell lysis), then 40 cycles of 94°C for 30 seconds, 58°C for 2.5 minutes, 68°C for 2.5 minutes, ending with a melt curve. A positive result was Ct<40 and peak Tm of 77-79°C. A positive result for both targets was interpreted as carriage-positive, a negative result for both targets was interpreted as carriage-negative. Non-concordant results (positive/negative or negative/positive) were treated as a separate group. This assessment of carriage prevalence may be affected by several factors: recent antibiotic consumption, low microbial biomass, age under 6 months (as inferred from previous work [1]), or technical error during swab collection may lead to a lower estimate. Presence of dead bacterial cells in the nasopharynx or cross-contamination during sample handling may lead to false positives. The sample size of 100 was selected as adequate to encompass a predicted prevalence of 10-90% with a precision of 5% and 95% CI [35]. This sample size is approximately one tenth of the total population being estimated, i.e. all 12 month old children born in Maela between 2007-2010.

### Microscopy

A suspension of NPS STGG sample was fixed overnight at 4°C in 3% paraformaldehyde, and dehydrated in suspension with 96% ethanol. Fluorescent hybridisation with an Alexa546 labelled probe (Invitrogen) was performed on fixed sample in buffered suspension (20mM Tris-HCl, 0.9M NaCl, 0.1% SDS) for 2 hours at 55°C, and washed with 20mM Tris-HCl, 0.9M NaCl for 5 minutes at 55°C. The samples were then suspended in water and applied to standard microscope slides, dried, and a coverslip applied with Vectashield mounting medium (Vector Laboratories, California, USA). The probe with sequence GUUCUUCACCCCGAAAACG targets the V5 region of the 16S ribosomal RNA, corresponding approximately to position 822-840 of the 16S rRNA gene of *E. coli.* It was not found to bind to an ORT sample when processed in parallel. A Zeiss Axiovert 200M fluorescence microscope with Zeiss filter set 43 (Ex 545/25, FT 570, Em 605/70) was used to visualise probed cells. A Sony DXC-390p colour video camera was used for imaging, which was recorded using Debut video capture (v.4.04, NCH Software). Images were collected from captured video files using Camtasia video processing software (v.8.6, TechSmith).

### DNA extraction and sequencing

Preliminary work was performed on DNA extracted from swabs with FastDNA Spin Kit For Soil (MP Biomedicals, Ohio, USA) which was then amplified using multiple displacement amplification (MDA). MDA reagents were filtered at 0.2μm, endonuclease digested with ϕ29 enzyme, and UV irradiated (254nm) prior to use to remove any exogenous DNA from the subsequent amplification reaction [36]. 3μl of sample was heat denatured with 2μl Heat Denaturation Buffer (20mM Tris HCl pH 8.0, 2mM EDTA, and 400μM PTO random hexamers (Eurofins Genomics)) at 95°C for 3 minutes. 15μl of Reaction Master Mix (1x RepliPHI reaction buffer, RepliPHI ϕ29 enzyme (6.7 U/μl) (Epicentre), 0.5mM dNTP, 50μM PTO random hexamers (Eurofins MWG), 5% DMSO, 10mM DTT) was then added, samples were incubated at 30°C for 16 hours, and the reaction halted by final incubation at 65°C for 20 minutes. Eight separate reaction volumes were processed per sample, these parallel reactions were pooled before sequencing using the Miseq 250bp paired end protocol with a 450bp library fragment size.

### Genome analysis

Samples were selected to maximise recovery of the genome of interest by choosing those with a high proportional abundance of the bacterium from previous 16S rRNA gene sequencing data [1]. Raw reads from MDA-amplified samples were first classified using Kraken v. 0.10.6 [37] and parsed to remove any reads classified as mammal, *Moraxella, Haemophilus* or *Streptococcus.* The remaining reads were then assembled using SPAdes v. 3.10.0 [38]. Contigs shorter than 500bp or with average coverage below 4X were discarded. For the two well-assembled samples, a BLAST+ v. 2.7.0 [39] screen of all contigs against the nr database was used to discard those that closely matched other known nasopharyngeal bacteria. The contigs first brought forward for the draft genomes were those that had consistent, low-identity matches to ORT. Samples were then reciprocally compared using BLAST+ to find further contigs present in all runs. These curated contig sets were manually improved using Gap5 v. 1.2.14 [40] and targeted PCR for gap closure, resulting in two syntenic draft genomes. The other assemblies with large numbers of short contigs were screened by comparison against the draft genomes using BLAST+ and extracting all contigs with >10% length hit. Automated annotation of curated contigs was performed using Prokka v. 1.11 [41] and the RefSeq database [42]. ANI and AAI were calculated using the enveomics calculators [43, 44]. ANI used a minimum length 700bp and minimum identity 70%, with 1000bp window size and 200bp step. AAI used a 20% identity cut-off. POCP was calculated as described by Qin *et al* [8]. The core and accessory genomes were calculated using Roary v. 3.11.3 [9]. Phylogenetic trees were built using RAxML v. 8.2.8 [45].

### Protein modelling

The OH ToxA protein sequence was searched using the program FUGUE [46] against a database of all chains of the Protein Data Bank (PDB) as of June 2017 [47]. Significant similarity with 30% sequence identity was found for residues 554-1269 to chain X of PDB ID: 2EBF [48], with corresponds to the C-terminal region of the *Pasteurella* mitogenic toxin. The matched region was aligned to the chain sequence using FUGUE, and models were generated with MODELLER v.9.15 using “very slow” refinement [49]. Visualization of the resulting models was performed on PyMOL Molecular Graphics System v1.8.

### Culture methods

NPS STGG samples were streaked out on blood or chocolate agar and incubated at 37°C in aerobic, enriched CO_2_ or anaerobic conditions for 48 hours. 25ml brain heart infusion (BHI) broths, with and without 1μg/ml ampicillin, were inoculated with 5ul of STGG and incubated either static or shaking at 37°C for up to one week. Other nasopharyngeal species were recovered from NPS STGG following these methods, but OH was not.

### Accession numbers

Accession numbers for the 7 samples are listed in Table 1, and are deposited under project accession ERP107699 (ERS2321170-6). The ORT genomes used for comparison are UMN-88, accession CP006828.1 and DSM 15997, accession CP003283.1.

## ACKNOWLEDGEMENTS

The authors acknowledge the generosity of Paul Wigley and Rachel Gilroy at the University of Liverpool for provision of an ORT strain as a negative control for fluorescent probe testing, the support of the SMRU laboratory and clinical teams, the Sanger Institute Pathogen Informatics team and the core sequencing and informatics teams.

